# Linkage-based ortholog refinement in bacterial pangenomes with CLARC

**DOI:** 10.1101/2024.12.18.629228

**Authors:** Indra González Ojeda, Samantha G. Palace, Pamela P. Martinez, Taj Azarian, Lindsay R. Grant, Laura L. Hammitt, William P. Hanage, Marc Lipsitch

## Abstract

Bacterial genomes exhibit significant variation in gene content and sequence identity. Pangenome analyses explore this diversity by classifying genes into core and accessory clusters of orthologous groups (COGs). However, strict sequence identity cutoffs can misclassify divergent alleles as different genes, inflating accessory gene counts. CLARC (Connected Linkage and Alignment Redefinition of COGs) [https://github.com/IndraGonz/CLARC] improves pangenome analyses by condensing accessory COGs using functional annotation and linkage information. Through this approach, orthologous groups are consolidated into more practical units of selection. Analyzing 8,000+ *Streptococcus pneumoniae* genomes, CLARC reduced accessory gene estimates by more than 30% and improved evolutionary predictions based on accessory gene frequencies. By refining COG definitions, CLARC offers critical insights into bacterial evolution, aiding genetic studies across diverse populations.

## Background

Advances in genomic sequencing technologies have highlighted the significant genetic diversity that can exist within a bacterial species^(1–3)^. Bacteria accumulate variation in both sequence identity and gene content through vertical transmission of genetic material and multiple means of horizontal gene transfer^(4)^. These genetic differences can underlie key phenotypic traits, and understanding them can aid in the study of clinically relevant qualities such as antimicrobial resistance, virulence, and vaccine susceptibility^(5–7)^, alongside other ecologically important features of bacterial evolution. The need to accurately represent a population’s diversity prompted the conceptualization of the bacterial pangenome, defined as the entire set of genes among all members of a species^(8)^.

Within the pangenome, genes can fall in one of two categories: *core genes* which are those present in all (or most) members of the population or *accessory genes* which are genes present in only a subset of the population; these two categories offer unique and complementary information. For example, accessory genes have played a key role in evolutionary models of bacterial population structure and dynamics, helping to understand the maintenance of genetic diversity^(9)^ and estimating gene gain and loss rates^(10)^. However, to maximize insights into the bacterial pangenome, genes within a population must first be accurately predicted.

Workflows to define the pangenome of a population generally cluster annotated sequences into orthologous groups based on sequence homology^(11–15)^. This produces clusters of orthologous genes (COGs), which are considered to represent distinct genes in the population^(16)^. Choosing a homology cutoff to generate COGs produces a trade-off between specificity and sensitivity; a high sequence identity cutoff decreases the chance of clustering distantly related sequences, but it also may exclude divergent alleles of the same homologous locus, and vice versa. This becomes especially important in genetically diverse species, where sacrificing sensitivity can inflate accessory gene estimates by misclassifying alleles of the same gene into separate COGs. Indeed, recent work in different pathogens highlights how annotation errors and sequence identity cutoffs can affect pangenome estimates^(17–19)^ with a significant impact on downstream analyses. This is particularly true for analyses that rely on gene frequency estimates such as studies of gene dynamics across populations or time^(20, 21)^, or association studies (e.g., pangenome-wide association studies, panGWAS)^(22)^.

Here we propose a new approach that combines sequence identity with functional and linkage information to correct for the over-splitting of gene variants into multiple clusters. This algorithm has been packaged into a bioinformatics tool named **CLARC** (**C**onnected **L**inkage and **A**lignment **R**edefinition of **C**OGs). CLARC takes the outputs of current pangenome tools as the input to generate refined COG definitions. We leverage the population-dependent definition of accessory genes to identify and cluster accessory COGs that correspond to ‘similar units’ and then condense them. These clusters are created by identifying COGs that never co-occur in a given population of samples, that also share sequence identity and functional annotation. Under this framework, COGs that perfectly exclude each other and have similar functional annotation might be variants of the same gene, or genes of a different origin that perform a homologous function, and clustering these can result in more evolutionarily informed gene boundaries.

We have tested CLARC by refining the COG definitions in pangenome analyses of *Streptococcus pneumoniae*, a species that undergoes homologous recombination through natural transformation and is characterized by variation in genome content. We demonstrate that CLARC results in improved accessory and core gene determination and that the re-defined COGs may represent a better measure of the practical units of selection in this bacterial species.

## Results

### Developing a bioinformatics tool to refine COG definitions in bacterial pangenomes

CLARC uses a custom clustering algorithm to identify and reduce redundancy in COG definitions (Figure 1A). The first step in this process identifies accessory COG pairs that appear to be the ‘same unit’ using 3 constraints: (1) COGs never co-occur in the same isolate, and (2) COGs meet a custom cutoff of nucleotide sequence similarity and (3) COGs get classified into the same functional group.

**Fig 1.**
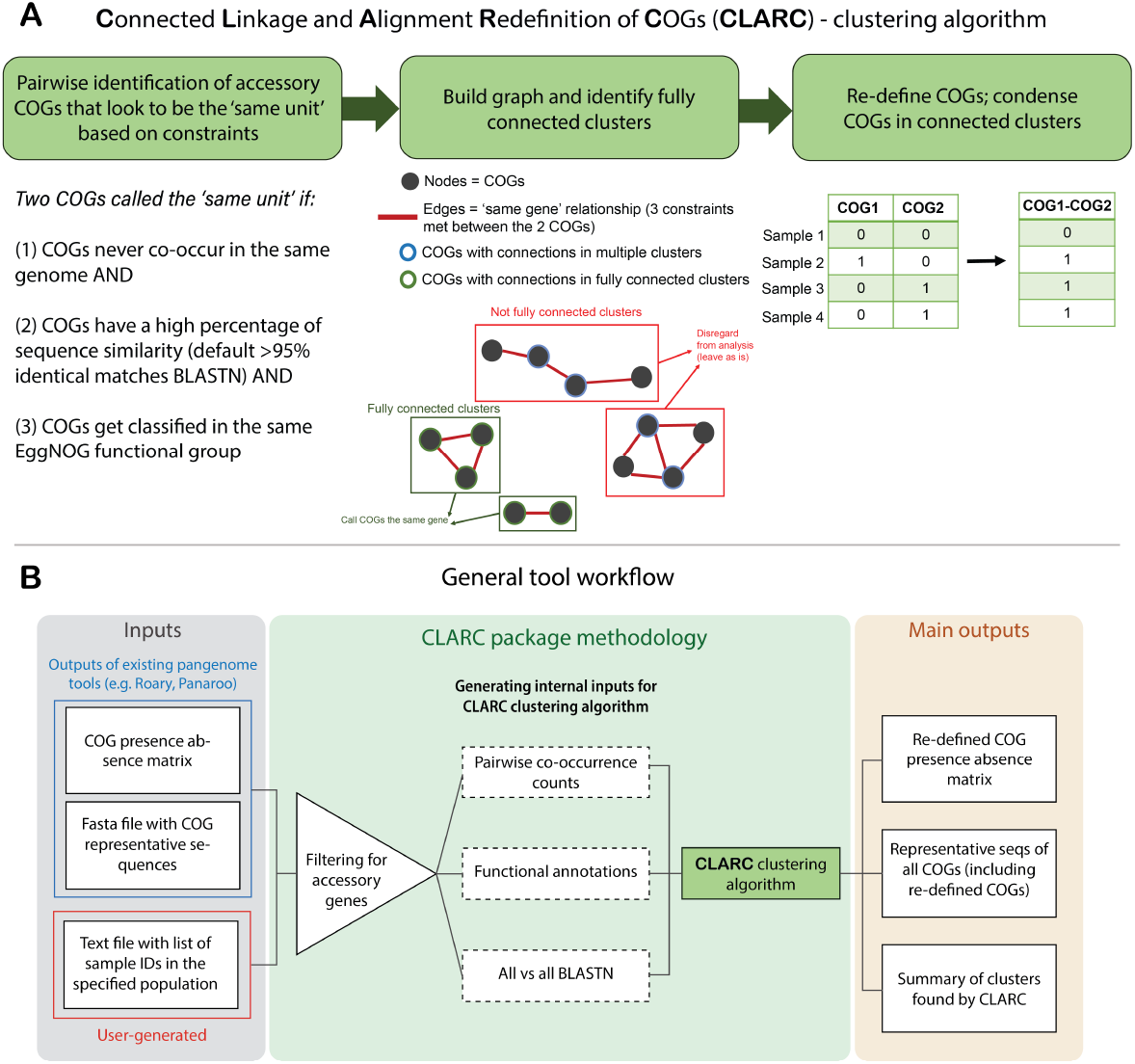
Workflow of CLARC tool. **(A)** Clustering algorithm used to reduce redundant COG definitions. Algorithm identifies ‘same unit’ clusters of 2 or more COGs based on sequence homology, mutual exclusivity across isolates and the same functional classification. COGs in each ‘same unit’ cluster are condensed into a re-defined COG. **(B)** Workflow of the CLARC bioinformatics package, including the clustering algorithm. When using the tool, the user inputs a csv file with the COG-presence-absence matrix previously generated by a current pangenome tool, the fasta file containing the COG representative sequences for that pangenome analysis, and a text file outlining the samples across which linkage will be evaluated. CLARC outputs the re-defined COG classifications in the same format as current tools, to facilitate downstream analyses.

Secondly, the algorithm builds a graph where each node represents a COG, and two nodes are connected by an edge if all constraints were met for that COG pair in the previous step. The algorithm then looks for fully connected clusters of COGs that appear to be the same unit. By fully connected, we mean clusters where all COGs have connections to all other COGs in the cluster. This ensures that all COGs within the cluster are mutually exclusive and helps prevent false positives from low frequency genes. COGs that are not in fully connected clusters within the graph are not modified further. Finally, the pipeline condenses COGs within these ‘same unit’ clusters by summing their individual presence absence matrices.

The clustering algorithm is implemented in the CLARC bioinformatic tool (Figure 1B). This tool takes the COG classifications of current pangenome tools, calculates all internal inputs, and outputs the re-defined COGs created by the clustering algorithm. The current version of CLARC (v.1.1.0) supports raw inputs from Roary^(11)^ and Panaroo^(15)^. However, the results from any pangenome analysis can be used if the inputs are formatted like the results from these tools. In addition to the output files of current pangenome tools, CLARC requires a text file with a sample ID list for the genomes that compose the population of interest. Linkage will be calculated based on this population, which can be all samples or a subset of samples if a distinct population within the whole set is known (e.g., samples collected in a specific geographic location).

Instructions on how to install and use CLARC, as well as descriptions of additional output files and other tool options can be found at https://github.com/IndraGonz/CLARC.

### Large collection of genetically diverse S. pneumoniae genomes worldwide to illustrate impacts of pangenome refinement

Including more samples in a pangenome analysis improves its accuracy in capturing a species’ full genetic diversity. *S. pneumoniae* is typically carried asymptomatically but can cause pneumonia, meningitis, and sepsis in a small proportion of cases, contributing to a significant disease burden. There has been extensive sampling coupled to whole genome sequencing of pneumococcal carriage worldwide, spearheaded by the global pneumococcal sequencing project (GPS) (https://www.pneumogen.net/gps/). This deep sampling in *S. pneumoniae* provides an opportunity to test CLARC on large-scale pangenome analyses (thousands of genomes) in this genetically diverse bacterial species.

We selected 7 datasets collected from nasopharyngeal carriage samples across 4 different continents, totaling 8,898 genomes (Figure 2A and 2B, see methods section for full description of the datasets). To evaluate the genetic diversity across all datasets we classified samples into lineages using an established k-mer based method, where genomes are assigned into different Global Pneumococcal Sequence Clusters (GPSCs)^(23)^. We observe clear geographic signatures in the distribution of GPSCs, with different locations having different clonal compositions (Figure 2C). When calculating Mash^(24)^ distances across all samples, we find that the genomes have an average genomic distance of 1.24% (**Supplementary figure S1**). These results suggest that our genome collection captures the diversity within *S. pneumoniae*.

**Fig 2.**
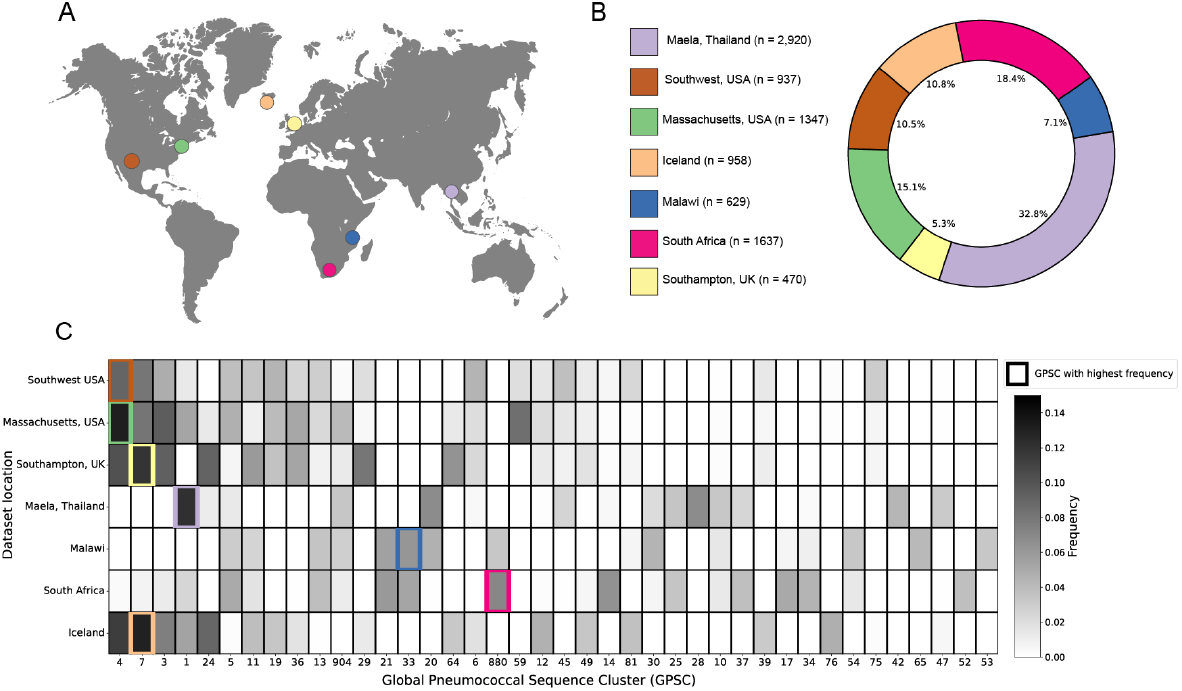
Genetic composition of pneumococcal carriage datasets from diverse populations worldwide. **(A)** Map showing the geographic location where samples were collected in each dataset. **(B)** Detailed breakdown of the number and proportion of samples from each carriage dataset in the full set of genomes. **(C)** Distribution of most prevalent global pneumococcal sequence clusters in the different populations. Only the 40 most common GPSCs across all datasets are displayed. The most common GPSC in each population is highlighted with a colored outline.

### CLARC consistently reduces gene oversplitting and increases accuracy of core genome determination in pneumococcal pangenome analyses

To evaluate the accumulation of core and accessory genes in *S. pneumoniae* as genomes are added, we generated 7 different pangenome analyses by sequentially adding each of the carriage datasets one by one and then counting the number of core and accessory genes. We used Roary ^(11)^ to define the clusters of orthologous genes (COGs) present across samples, since it is a widely used and computationally efficient tool.

Since the *S. pneumoniae* pangenome is on the ‘extreme’ end of an open pangenome, we expect the core genome to decrease in size and the accessory genome to increase in size as samples are added into the analysis. Theoretically, the number of genes found per new genome decreases logarithmically^(25)^. For *S. pneumoniae*, previous research estimates that after 100 genomes the number of new genes found per added genome should be less than 10^(26)^. Thus, after around 1,000 genomes we would expect the core and accessory gene counts to stabilize. Instead, we observe a sharp decrease in the number of core genes, coupled to a dramatic increase in accessory genes as datasets are added (Figure 3A and 3B, solid lines). The sharpest changes occur when adding the samples from Maela, Thailand into the pangenome analysis. This might be due to the fact that it is the largest dataset (2,920 genomes) and also the most genetically distinct, as seen in Figure 2C.

**Fig 3.**
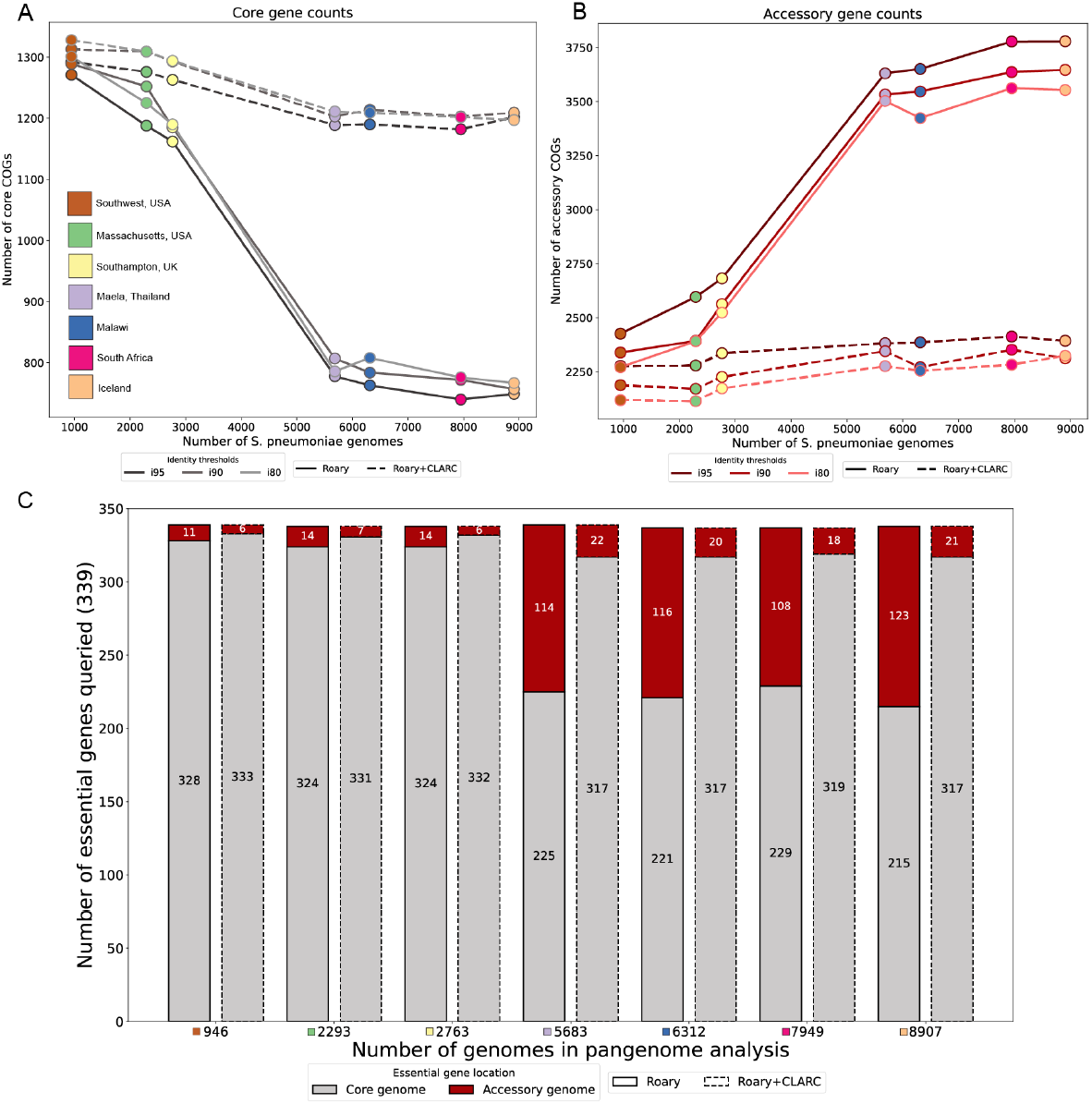
Effect of CLARC on the accumulation of core and accessory genes and the misclassification of essential genes as datasets are added to the pangenome analysis. **(A)** and **(B)** core and accessory gene counts as datasets are added into the analysis, respectively. Here, core genes were defined as those present in >95% of samples, and accessory genes as those present in 5–95% of samples within a single target population (Southwest US). Results are consistent across various Roary sequence identity thresholds (95%, 90% and 80%). **(C)** Query of essential genes in the pneumococcal pangenome analyses. The count of essential genes found in the core genome is shown in gray, and those misclassified as accessory are shown in red. In the largest pangenome analysis. Roary+CLARC refinement is shown to consistently correct the misclassification of essential genes as accessory across all analyses.

These results deviate from expectation, suggesting that the accessory genome might be inflated at the expense of the core genome. This is consistent with previous research that has reported the inflation of the accessory genome and/or the underestimation of the core genome for different bacterial species^(17, 18)^.

To test this, we leverage gene essentiality. Essential genes are necessary for an organism’s survival, so they are a part of the core genome. We acknowledge that essentiality can be dependent on strain background and environmental conditions^(27)^, but an accurate pangenome analysis should still identify a significant proportion of essential genes as core. Previous work by van Opijnen et al. (2009)^(28)^ characterized a set of 339 essential genes in *S. pneumoniae* using Tn-Seq. We extracted the sequences of these genes and searched for them in the core and accessory genomes identified in each of the previous pangenome analyses. In the pangenome built with all samples, 36% of essential genes were missing from the core genome and found in the accessory genome (Figure 3C, bars with solid outline). This is consistent with core genes being misclassified as accessory, inflating the accessory genome.

CLARC refinement of all pangenome analyses stabilized core and accessory gene counts (Figure 3A and 3B, dashed lines) and dramatically reduced the number of essential genes misclassified as accessory (Figure 3C, bars with dashed outline). This suggests that CLARC successfully increases the accuracy of COG definitions in the *S. pneumoniae* pangenome.

### Clusters found by CLARC dramatically reduce the accessory genome and are enriched in functions commonly found in core genes

We dive deeper into the clusters identified by CLARC using the pneumococcal pangenome built from all samples. Here, CLARC found 727 clusters that included 1,660 COGs. After condensing these clusters, the majority of the redefined COGs were reclassified as part of the core genome; this reduced the accessory genome by 37% (Figure 4A). Most of these clusters were composed of COG pairs, with the largest clusters containing 6 COGs (Figure 4B).

**Fig 4.**
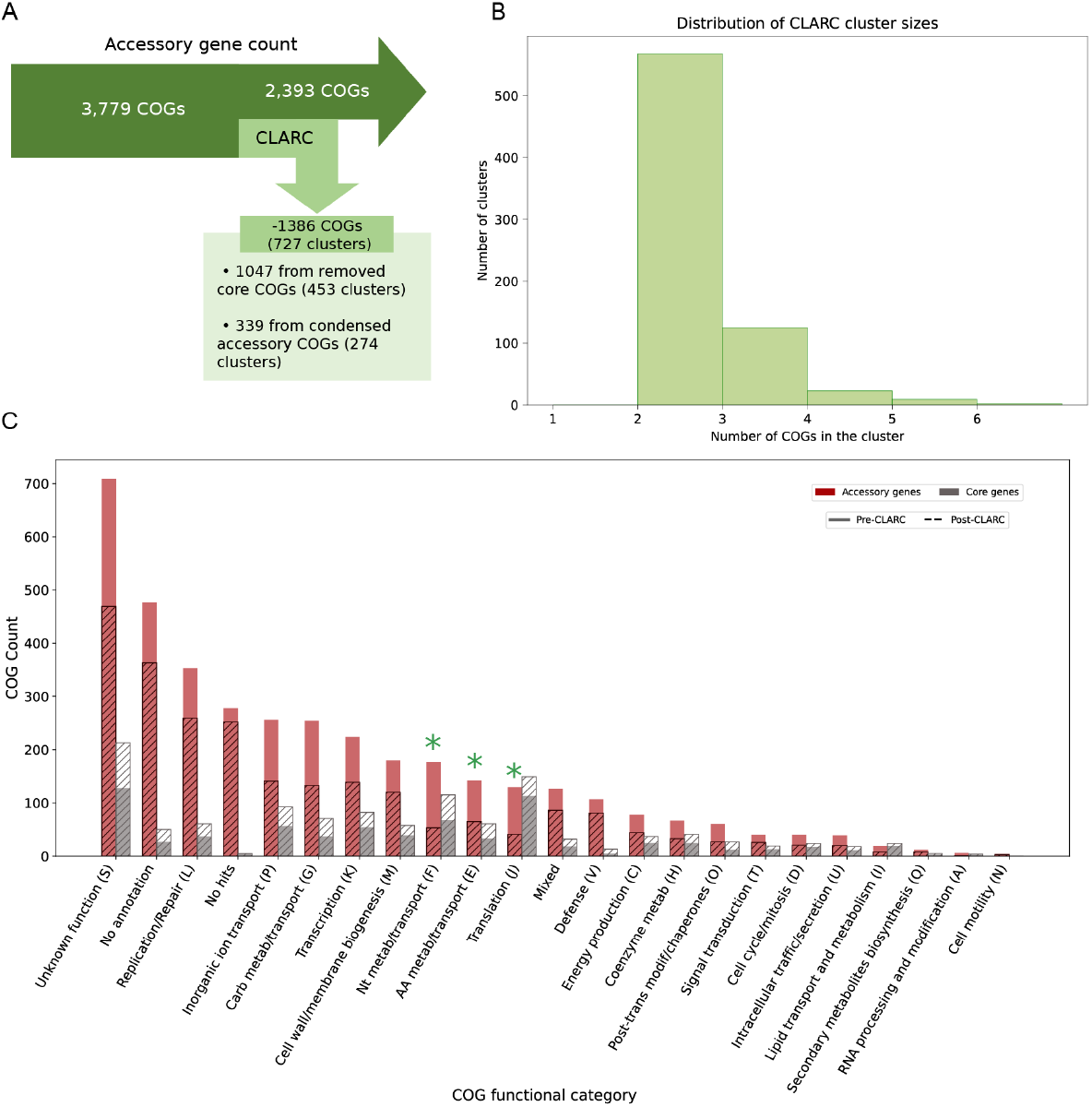
Analysis of CLARC clusters found using a pangenome analysis generated from all carriage datasets. The pangenome analysis used here was built using Roary default parameters. We focus this targeted analysis on the Southwest USA population to evaluate the impact CLARC has on the accessory genome of a single closed population. **(A)** Breakdown of clusters found by CLARC and reduction of accessory genome. **(B)** Distribution of the number of COGs forming the clusters found by CLARC. **(C)** Distribution of COG functional groups in the accessory and core genomes before and after CLARC refinement.

COGs identified in CLARC clusters were spread across the full range of functional categories (Figure 4C). The most significant changes after CLARC refinement occurred in the categories of nucleotide metabolism/transport (F), amino acid metabolism/transport (E), and translation (J), with their presence in the accessory genome reduced by over 50% (highlighted with green asterisks). These categories represent important cellular functions and are commonly enriched in the core genome of various bacterial species^(29)^.

### Refined gene definitions improve the prediction of S. pneumoniae post-vaccine population structure

Because CLARC uses linkage and functional information to re-define COGs, we wanted to test if the new gene definitions represent a more accurate unit of selection in *S. pneumoniae* populations. For this, we leverage previous work that describes how accessory genes in *S. pneumoniae* respond to vaccine perturbation^(30, 31)^.

The pneumococcal conjugate vaccine (PCV) was deployed worldwide in the early 2000s to decrease the public health burden of pneumococcal disease^(32)^. This vaccine targets some S. pneumoniae serotypes, creating genetic perturbations in the population. Corander et al. (2017)^(30)^ found that after vaccination, strain distributions shifted (targeted strains decreased in frequency), but accessory gene frequencies remained stable, even across geographically distinct populations with differing strain compositions. This stability is attributed to negative frequency-dependent selection (NFDS) acting on the accessory genome, where variant fitness is inversely correlated with abundance^(33)^. Thus, low frequency variants will have a competitive advantage, and traits under NFDS will be driven to an intermediate frequency equilibrium. Since this study follows accessory gene frequencies, inaccurate gene frequency estimates could affect the results. Using the samples from the Southwest US population, we compared the pre-vaccine vs post-vaccine accessory gene frequencies before and after correcting through CLARC (Figure 5A and 5B). COGs forming CLARC clusters were located through the whole frequency range, with a higher density around low frequency genes. This makes sense because the over-splitting of genes will cause an inflation of lower frequency genes. Nevertheless, accessory gene frequencies are still conserved before and after vaccination in the post-CLARC gene definitions, even when the number of accessory genes is greatly reduced.

**Fig 5.**
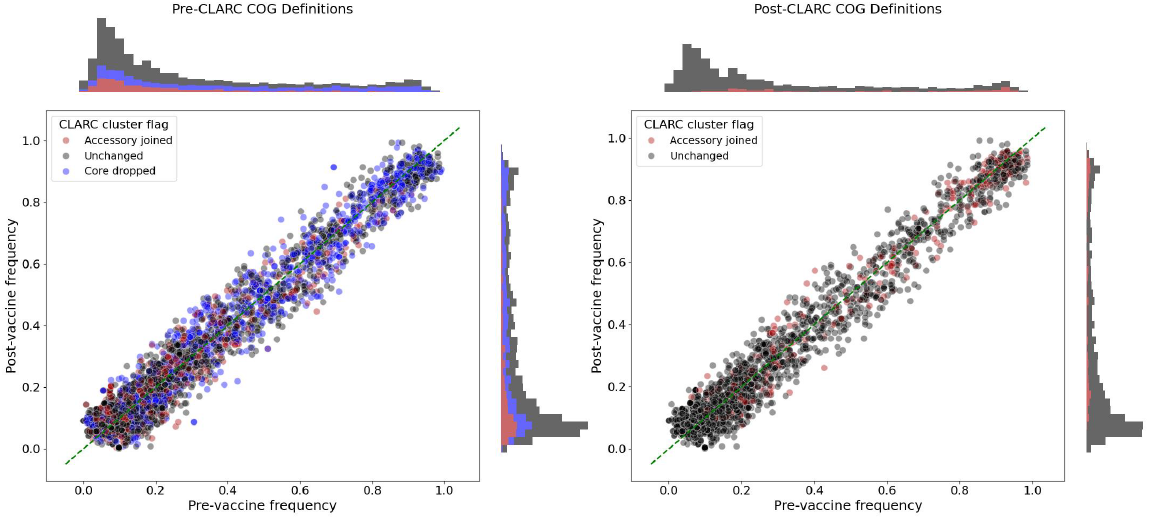
Effect of CLARC in the pre- and post-vaccination frequency of accessory genes in the Southwest US population. **(A)** Using pre-CLARC gene definitions and **(B)** using post-CLARC accessory gene definitions. COGs identified as part of a CLARC cluster are colored. Blue dots represent COGs identified as part of a cluster that turned out to be a core gene when condensed, while red dots represent COGs that were part of a cluster that remained an accessory gene after CLARC correction. In (B) the blue dots don’t appear because they were eliminated from the set of accessory genes (since they were re-classified as core genes), but the new COGs representing condensed accessory gene clusters are marked in red. The frequencies plotted for the post-CLARC genes are the condensed frequencies of these new COGs.

If accessory genes are under balancing selection, the population will respond to the vaccine perturbation by favoring strains that can best restore the pre-vaccine accessory gene frequencies. Azarian & Martinez et al. (2020)^(31)^ demonstrated that the post-vaccination population structure (GPSC composition) of *S. pneumoniae* could be predicted using a quadratic programming model. This model restored the prevaccination equilibriums of accessory genes using the strains not targeted by the vaccine (see methods section for detailed information on this model). The predictive power of this model relies on having accurate accessory gene definitions that represent units under balancing selection. If a gene (or a particular function under selection) is split, then the model will try to restore the inaccurate frequencies instead of their actual equilibrium frequencies, resulting in a less accurate prediction. Therefore, we can use the accuracy of this prediction as a proxy for the quality of pangenome COG classifications.

We predict the post-vaccination GPSC frequencies in the Southwest US dataset using the accessory gene frequencies of a Roary pangenome analysis (Figure 6A) and the accessory gene frequencies after running CLARC (Figure 6B). We find that using the CLARC re-defined COGs slightly improves the prediction, even when fewer accessory genes are given to the model (2,393 accessory genes in the Roary-CLARC prediction vs 3,779 in the Roary prediction).

**Fig 6.**
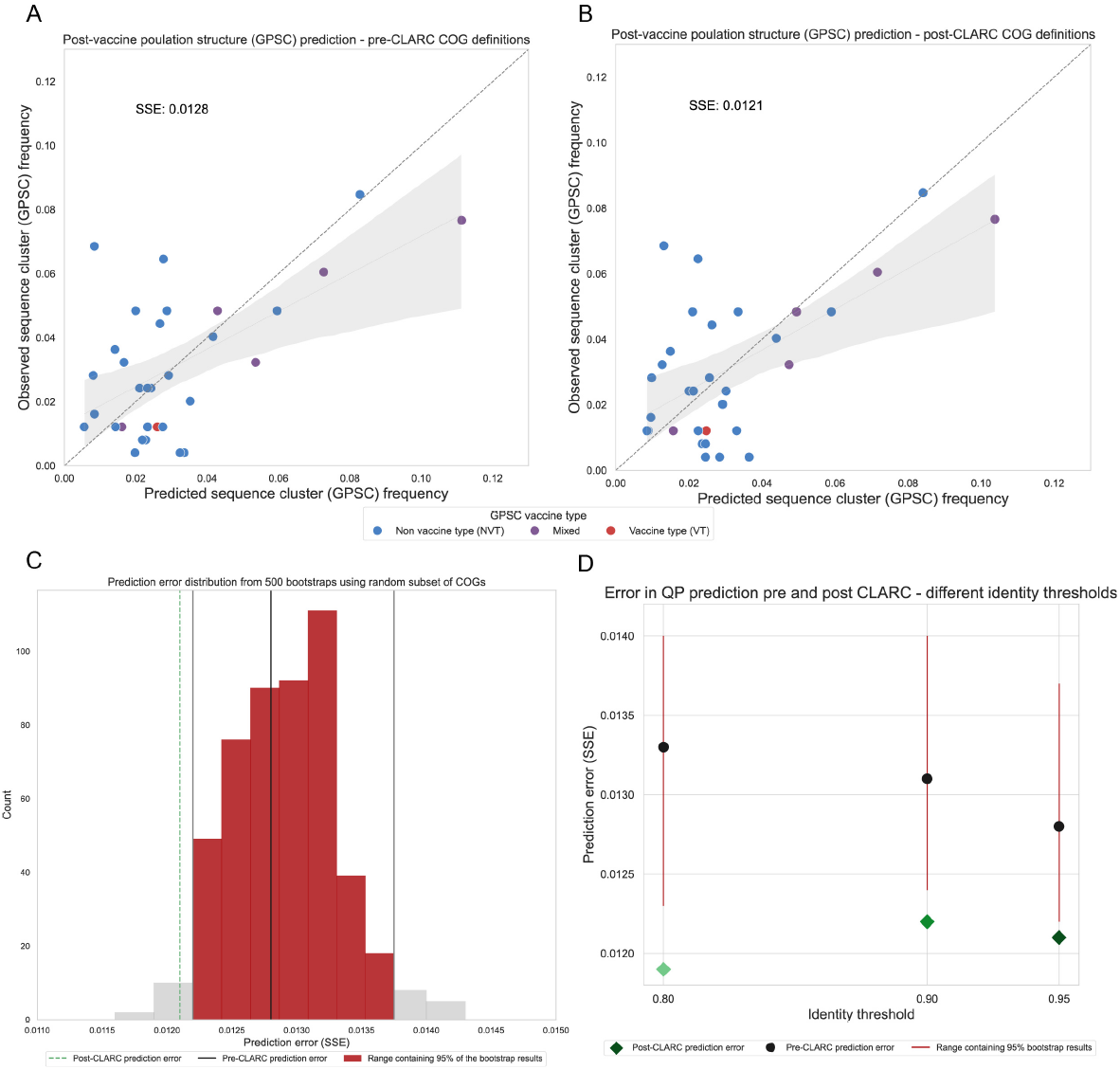
Effect of CLARC on the prediction of pneumococcal post-vaccine population structure. The COG definitions for these predictions come from the all-carriage pangenome. **(A)** Population structure prediction in the Southwest US using the accessory genes definitions generated by Roary. Prediction is compared to the observed post-vaccine GPSC frequencies in the population. Sum of squared errors is used to evaluate the prediction. **(B)** Population structure prediction in the Southwest US using the accessory genes definitions after CLARC refinement. **(C)** Comparison of prediction improvement against random bootstraps with a subset of the original Roary accessory COGs. This is for the all-carriage pangenome run on default parameters. **(D)** Improvement of model prediction with CLARC refinement, using various Roary identity threshold parameters.

However, this difference in prediction error might result from a smaller subset of accessory genes coincidentally improving the prediction. To test whether the post-CLARC improvement was truly due to the redefined COG frequencies, we performed a bootstrap analysis. We randomly selected 2,393 genes from the original 3,779 Roary accessory gene list, used them for prediction, and recorded the resulting prediction errors (see histogram in Figure 6C). The lower error achieved with the CLARC gene set falls outside of the 95% prediction interval of the random bootstraps, and this prediction improvement is independent of the identity threshold used to run Roary (Figure 6D). Taken together, these results are consistent with CLARC providing gene definitions that represent a practical unit of selection in this system.

#### Comparison with graph-based pangenome tool

Pangenome tools like Roary^(11)^, PanX^(12)^, PIRATE^(13)^ and various others use protein clustering algorithms like CD-HIT^(34)^ or MCL^(35)^ to cluster annotated protein sequences into orthologous groups, without using gene neighborhood information to generate the clusters. On the other hand, recent tools such as Panaroo^(15)^ and Ppanggolin^(14)^ build genome graphs to identify orthologous clusters. Prioritizing spatial information can help mitigate the effects of sequencing errors and incorrect annotations. However, if the synteny of genes within a species is not consistently conserved, this approach may lead to misclassifications.

To test the impact of CLARC on results from a graph-based pangenome method, we used the different *S. pneumoniae* datasets (as seen in Figure 2) to generate pangenome analyses using Panaroo. CLARC did identify a handful of clusters on many of the Panaroo pangenome results (**Supplementary tables S1-S2**). Nevertheless, the clusters found in Panaroo COG definitions were orders of magnitude fewer than in

Roary analyses. However, using the accessory genes identified with Panaroo in the NFDS population structure prediction model consistently resulted in significantly higher error than those generated with Roary (Figure 7). This was true across different parameters and identity thresholds. For both Panaroo and Roary, re-defining the accessory genome with CLARC generally resulted in improved population structure predictions. The results suggest that using the pangenome generated with Roary+CLARC provides the highest resolution to study negative frequency dependent selection in the accessory genome of *S. pneumoniae*.

**Fig 7.**
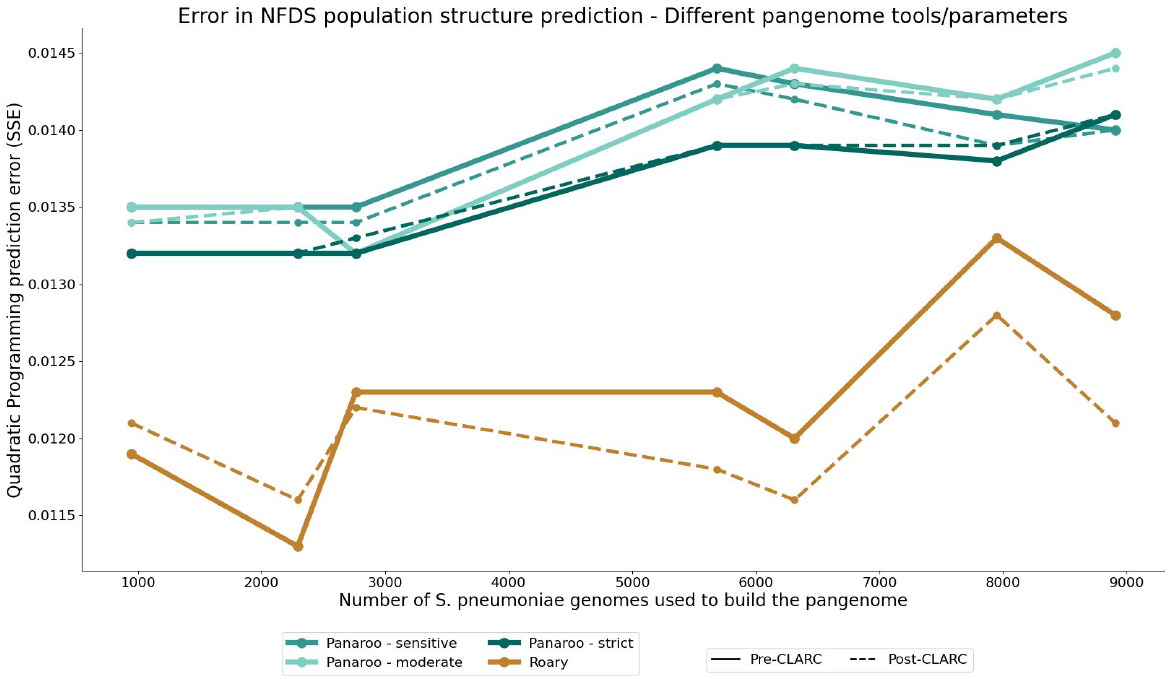
Performance of accessory gene definitions from different tools in the S. pneumoniae population structure prediction model. Pangenome analyses were built using their respective default parameters.

## Discussion

Bacterial pangenome analyses aggregate coding sequences into clusters of orthologous groups (COGs), to identify genes shared across members of a population. These COGs are meant to highlight shared evolutionary pressures and genetic diversity across strains. Current pangenome tools broadly use sequence similarity to generate clusters, then identify paralogs using synteny-based^(15, 36)^ or tree-based approaches^(12)^, among others. However, functional classification information is generally not used to generate the COG definitions. Albeit sequence content is often correlated with function, it is not always fully predictive^(37)^; diverged sequences can perform similar functions due to structural or functional convergence, while nearly identical sequences can evolve distinct roles under different selective pressures^(38)^.

Current methods can erroneously split functionally related genes into different orthologous groups, or cluster genes with diverged functions into the same group. Therefore, there is a growing push to adopt a more functional approach to pangenomic gene classification^(39, 40)^. Ultimately, selection acts on function (and not necessarily sequence), so having a functionally informed classification of COGs could optimize evolutionary studies involving pangenomes. This approach could be particularly valuable for reducing the accessory gene set in bacteria with open pangenomes, helping to mechanistically link specific genes to observed traits or phenomena.

To address these gaps, we have developed CLARC, a bioinformatics package that refines the core and accessory gene definitions of existing pangenome tools. This package is computationally efficient and can be run locally on pangenome analyses that include thousands of genomes. It can generate condensed COG definitions in just a few hours, making it practical for large-scale studies (**Supplementary figure S2**).

Our results indicate that Roary, a widely used pangenome tool, inflates the accessory genome of *S. pneumoniae* largely by splitting core genes into multiple accessory COGs. The prevalence of these misclassifications increases as genomic diversity is added into the pangenome analysis. CLARC addresses this inaccurate clustering, reducing the size of the pneumococcal accessory genome by over 30%. Additionally, the COG definitions generated with CLARC perform significantly better in a negative frequency dependent selection-based model that predicts the population level evolution of *S. pneumoniae* using accessory gene frequencies. Suggesting that CLARC provides COG definitions that function as more effective units of selection in this system. While we have tested CLARC using *S. pneumoniae* as a test case, there is no reason to believe the approach is limited to this organism. Just as the bioinformatic tools used to estimate the accessory genome may be used for many organisms, CLARC is capable of refining their results independently of the organism under study.

We are aware that Roary is part of the first generation of pangenome tools and thus it lacks quality control methods that more recent tools have implemented to correct for gene prediction errors. Most recently, graph-based pangenome tools like Panaroo have been shown to enhance gene definitions. When running CLARC on pangenome analyses created with Panaroo, some COG clusters are found but at a significantly lower rate than in the Roary analyses. However, the gene definitions generated with Roary+CLARC significantly outperform the Panaroo ones when studying negative frequency dependent selection in the accessory genome of *S. pneumoniae*. This highlights that CLARC produces clustering distinct from Panaroo, offering researchers a valuable new tool for obtaining functionally informed COG classifications.

Another important consideration is that graph-based methods are substantially more computationally intensive than other clustering methods^(15, 36)^. This makes them less tractable for analyses containing thousands of genomes, such as our pneumococcal carriage datasets. As more genomes become available, there is an increasing need for scalable clustering algorithms that can capture the diversity of the whole population. This is an important reason as to why Roary was used as the baseline to our analyses in lieu of a graph-based method. Running a simpler tool like Roary followed by CLARC resulted in accurate gene definitions, which provides a scalable alternative for researchers working with large genomic collections. Furthermore, it is possible that CLARC’s functionally informed approach may improve COG definitions at higher taxonomic levels (beyond the single species level), but further work is needed to confirm this.

With that said, there are some limitations to consider when using CLARC. For example, the linkage constraint in CLARC’s clustering algorithm does not account for multi-copy genes within a genome. If a genome contains two variants of the same gene classified as separate COGs, CLARC will not resolve this misclassification. Thus, CLARC may offer less benefit for genes that are frequently multicopy intrachromosomally. Additionally, CLARC runs efficiently under default parameters by restricting clustering to COGs present in at least 5% of the population, excluding singletons and very low-frequency COGs. While this threshold can be adjusted to include low-frequency genes, doing so would likely increase computation time significantly, as pairwise calculations grow non-linearly with the number of accessory genes included.

## Conclusions

Identifying genes shared across members of a bacterial population continues to be an ongoing challenge in bacterial genomics. As the number of sequenced genomes grows, so does the need for functionally informed gene classifications that can capture the diversity of gene functions and their evolutionary significance. Towards this goal we developed CLARC: a tool that combines functional classifications with linkage information to re-define COG classifications generated by current pangenome tools. CLARC seeks to create more evolutionarily informed gene boundaries by condensing redundant definitions of accessory COGs. Therefore, CLARC is meant to complement (and not replace) current tools.

CLARC is particularly useful for researchers conducting analyses that depend on COG frequencies, such as examining the evolutionary dynamics of individual genes. Its benefits are especially pronounced in large-scale pangenome studies that involve thousands of genomes, where it can efficiently clarify gene relationships. Overall, a next step in the analysis of bacterial pangenomes is to achieve greater accuracy in COG definitions as units of selection, and CLARC represents a practical advancement in this direction.

## Methods

### S. pneumoniae datasets used

In total, 8,907 pneumococcal genomes were used in this study. 8,898 genomes are from pneumococcal carriage datasets worldwide that have been indexed by the global pneumococcal sequencing project (GPS, https://www.pneumogen.net/gps/). These collections correspond to 7 distinct studies where a defined host population was systematically sampled, named after the geographic location in which they were collected: Massachusetts, Southwest USA, Maela, Southampton, Iceland, Malawi, and South Africa. The remaining 9 genomes used correspond to closed reference assemblies obtained from the NCBI database (https://www.ncbi.nlm.nih.gov/). The accession numbers for all samples can be found in the **Additional file 2** document.

The Massachusetts (MA), USA pneumococcal dataset consists of 1,347 isolates taken from the nasopharynxes of children of up to six years of age during routine primary care physician visits. These isolates were collected during two sampling periods, 2001-2007 (616 genomes)^(41)^ and 2009-2014 (731 genomes)^(42)^. The Southwest USA dataset totals 937 genomes which are a subset of 3 studies of pneumococcal carriage conducted among Native American communities from 1998 to 2012, as previously described^(43–45)^. The Maela collection is composed of 3,085 samples isolated from the nasopharynxes of infants of up to two years of age and their mothers, in a Thai refugee camp^(46)^. Of the Maela genomes, 2,920 passed the assembly quality control filter imposed during downstream analyses. The Southampton, United Kingdom collection includes 516 genomes isolated from the nasopharynxes of children of up to four years of age during outpatient visits^(47)^; in this dataset 470 genome assemblies passed quality control filtering. The Iceland dataset consists of 958 isolates collected from the nasopharynxes of healthy children^(48)^. The Malawi dataset is composed of 629 isolates collected in rural Karonga District from a cohort of mother-infant pairs and household members <16 years during 2009-2014^(49)^. Finally, the South Africa dataset contains 1,637 samples collected from nasopharyngeal swabs from children under 12 years old and their mothers, sampled in the Soweto region of South Africa^(50)^.

### Assembly bioinformatics pipeline

Short-read Illumina whole genome sequencing data was available for all isolates. Raw sequence reads were downloaded from the Sequence Read Archive (SRA) in the NCBI database. Assembly from shotgun sequencing reads was performed as previously described^(51)^. Genomic assembly from raw reads was constructed using SPAdes v3.10^(52)^ and annotated using Prokka v1.11^(53)^ embedded in Unicycler^(54)^. Quast v4.4^(55)^ was used to assess the quality of each genomic assembly. A sequence assembly was excluded when it had (i) N50 less than 15 kb; or (ii) ≥ 500 contigs, indicating the genome was too segmented; or (iii) a genome length *<* 1.9 Mb or *>* 2.4 Mb ; or (iv) a GPSC not assigned due to “genomes with outlying lengths detected.” with the reference genome ATCC 700669. All assemblies and annotated genomes generated are available in Zenodo at https://zenodo.org/records/14187853.

### Strain typing

To ensure consistency with worldwide efforts to study pneumococcus, each sample was typed for its global pneumococcal sequence cluster (GPSC), using a PopPUNK^(56)^ typing developed by the Global Pneumococcal Sequencing Project (GPS). This was achieved by inputting an external clustering file into PopPUNK, which contained information from all samples in the GPS database. The process followed the GPSC typing instructions provided by GPS. This approach resulted in nucleotide k-mer-based strain classifications with a global definition, allowing for comparison across different populations.

### Pangenome analyses

To evaluate the performance of different pangenome tools as pneumococcal genomes were added into the analysis, a series of seven cumulative pangenome analyses were conducted on the total of 8,907 genomes. In each analysis, a new dataset was incrementally added to those included in the previous analysis, beginning with a single dataset and culminating with all seven. Datasets were added in approximate order of geographic proximity, starting with samples from the Southwestern USA, followed by Massachusetts USA, Southampton UK, Maela Thailand, Malawi, South Africa, and Iceland. This cumulative approach enabled observation of the progressive impact of additional, geographically distinct datasets on the pangenome composition. All pangenome analyses included the 9 closed reference pneumococcal genomes. The breakdown of the sample IDs used in each analysis can be found in the **Additional file 2** document. To evaluate the tool specific differences of the analyses, each of the 7 distinct pangenomes were generated using both Roary v3.13.0^(11)^ and Panaroo v1.5.0^(15)^.

### Parameters in Roary pangenome analyses

Roary analyses were run using genome annotations as input, provided in GFF format. Each analysis was performed independently 3 times varying identity clustering thresholds (given by the -i flag): 98%, 95% identity (default), 90% identity and 80% identity. For all other options, default parameters were used. Intermediate files were kept using the -z flag.

### Parameters in Panaroo pangenome analyses

Due to the computationally intensive processes performed by Panaroo, it was not possible to run the larger pangenome analyses in one process. So, the first step in the analysis was to build pangenomes for each dataset individually. Then, all 7 of the aggregated pangenome analyses were built by merging the pangenomes of the corresponding datasets using Panaroo’s merge function. This was done using 4 different identity thresholds: 98% (default), 95%, 90% and 80%. Additionally, because Panaroo has three different cleanup modes (strict (default), moderate, and sensitive), analyses were run in each of these 3 modes. For all other options, default parameters were used. Scripts used to run each of the pangenome analyses can be found at https://github.com/IndraGonz/2024_GonzalezOjeda_CLARC.

### General description of CLARC pipeline workflow

The first step in the CLARC pipeline is to identify the set of accessory and core genes in the specified population. The default lower and upper bounds to filter for accessory genes are 5% and 95% respectively, and the lower bound to filter for core genes is 95%. COGs in between 5-95% of the population will be called accessory and COGs over 95% will be called core. When using the default parameters, CLARC does not consider genes in the cloud genome (genes only found in one or two isolates). This is because the cloud gene count can often be impacted by sequencing and assembly errors^(19)^, and linkage constraints become meaningless for COGs at very low frequencies. However, custom values for all these parameters can be specified by the user.

After the filtering step, CLARC uses the set of identified accessory genes to generate internal inputs for the clustering algorithm. This is done through scripts that generate the values needed to evaluate the 3 constraints used to cluster COGs. For the first constraint, a pairwise linkage metric is calculated by counting the number of genomes in which the accessory COG pairs co-occur (p_1_1) across samples in the specified population. The condition is met if p_1_1 = 0, since this means the COGs are never found in the same genome. For the sequence identity constraint, an all vs. all BLASTN^(57)^ alignment table is made by building a BLAST database using the COG representative sequences that are output from the original pangenome tool. Furthermore, for the final constraint, accessory COG sequences are functionally annotated into pre-established functional COG categories from the COG database^(58)^ using a custom script that queries each gene’s protein sequence against the EggNOG database v5.0^(59)^.

The generated inputs are fed into the clustering algorithm which identifies and condenses the ‘redundant’ COGs. CLARC then outputs the original presence-absence csv file, including the re-defined COGs. This csv file has the same format as output by the original pangenome tool. A fasta file with the representative sequences of all COGs (including redefined COGs) is also generated. When condensing the COG clusters, the sequence for the longest COG in the cluster is used as the representative sequence for that re-defined COG. These outputs are compatible with the original pangenome tools and thus the output from CLARC can be used as an input to any downstream analysis that takes input from Roary or Panaroo (e.g. Scoary^(60)^, etc.). A text file summarizing the CLARC clusters found is also created.

### Technical details of CLARC pipeline workflow

#### Initial filtering for core and accessory genes

An internal python script (filtering_acc_core.py) filters the original pangenome presence absence matrix to generate filtered presence absence csv files of the accessory and core genomes. This is done by calculating the frequency of each COG in the specified population of samples and then filtering for core and accessory genes using user-specified frequency thresholds. If run on the default parameters, CLARC will define accessory genes as those in between 5-95% frequency and core genes as those with greater than 95% frequency. The csv file with the accessory gene presence absence matrix in this step will be used to calculate the internal CLARC algorithm inputs.

#### Calculation of pairwise linkage of accessory genes

In this step, a python script (get_linkage_matrix.py) counts the different states of co-occurrence between every possible pair of accessory COGs. The possible states are: genomes where both COGs are observed together (P_11_), genomes where neither COG is observed (P_00_), or genomes where only one of the COGs is observed (P_10_ and P_01_). Thus, 4 of these matrices are generated, one per co-occurrence state.

#### Sequence identity comparison of accessory genes (all vs all BLASTN

A bash script (acccog_blastn.sh) uses all accessory COG representative sequences to build a BLASTN database and perform an all vs. all BLAST comparison. BLASTN version 2.15.0 is part of the internal conda environment created with the initial CLARC build during installation.

#### Functional annotation of COGs with the EggNOG database

CLARC uses a custom python script (eggnog_annotations.py) that serves as a wrapper of the EggNOG v.5.0 database^(59)^. Accessory COG representative sequences are translated into their protein sequences and submitted as queries to the database query interface. This allows for the remote functional annotation of COGs without having to download the EggNOG database locally (as is done when using tools like EggNOG Mapper^(61)^).

#### COG refinement with CLARC algorithm

Finally, a python script (clarc_condense.py) containing the CLARC clustering algorithm uses the internal inputs generated to condense the COG definitions and create the outputs for the re-defined COG list.

#### CLARC processing of pneumococcal pangenome analyses

CLARC v1.1.0 was run on the results of all pangenome analyses generated in this study. Raw outputs from Roary/Panaroo were used as input as specified in CLARC’s github repository. Additionally, the Southwest USA dataset was input as the population of samples used to calculate linkage. This population was used because it represents a closed population with extensive sampling before and after introduction of the 7-valent pneumococcal conjugate vaccine (PCV7). Default parameters were used in all analyses. Results for all CLARC runs can be found in this Zenodo repository.

#### Querying for pneumococcal essential genes

Previous research by van Opijnen et al.^(28)^ used Tn-Seq to identify 339 essential genes in a common strain of Streptococcus pneumoniae (TIGR4). The first step to query these essential genes in the core and accessory genomes defined by each pangenome analysis was to build a database containing the protein sequences of each of these essential genes, extracted from the TIGR4 genbank file of the strain used in the original analysis. Then, the protein sequences of the accessory and core genes in each pangenome analysis were compared to the database using BLASTP^(57)^ v2.14.1+. A hit was called if an alignment containing over 90% sequence identity and with an alignment length covering at least 90% of the coding sequence was found.

#### Quadratic programming prediction model

As described by Azarian & Martinez et al. (2020)^(31)^, the NFDS-based quadratic programming (QP) model uses the accessory gene composition of strains with a capsule type not targeted by the vaccine (non vaccine type, NVT) along with the prevaccine accessory gene frequencies in the population, to predict the post-vaccination strain frequencies. To test the effect of different accessory COG definitions on the performance of this prediction, we implement the model using the Southwest USA dataset as a test case. The extensive sampling before and after the introduction of the 7-valent pneumococcal conjugate vaccine (PCV7) in this dataset permits the comparison of the predicted post-vaccine population structure with the observed post-vaccine strain frequencies. In the United States the PCV7 was deployed in 2001, so we defined the prevaccine samples as those collected between 1998 and 2001 (274 genomes) and the post-vaccine samples as those collected between 2010 and 2012 (265 genomes). We focused on 32 GPSC strains that (1) had NVT taxa present before vaccination and (2) had more than 5 isolates in that strain cluster. For these strains, we computed the frequency of each strain such that their accessory gene frequencies would best approximate the pre-vaccine equilibriums using the quadratic programming approach. Quadratic programming involves optimizing a quadratic function based on several linearly constrained variables^(62)^ and was done using the package quad-prog v1.5.8 implemented in a jupyter notebook using the rpy2 python package with R v4.2.3. The quadratic programming algorithm was implemented within a jupyter notebook that calculated the pre-vaccine accessory gene composition of each strain (matrix containing the prevalence of each accessory gene in each GPSC under study) and the pre-vaccine global accessory gene frequencies (vector with the prevalence of each accessory gene in the pre-vaccine sample set) and input it into the quadratic programming function to obtain the predicted post-vaccine strain frequencies. The accuracy of the prediction was assessed using the sum of squared errors between the predicted and observed strain frequencies with the SSE function in the pracma R package v2.4.4. Different pangenome analyses were assessed by varying the accessory gene input matrix and recording the SSE of the corresponding prediction.

## Declarations

### ETHICS APPROVAL AND CONSENT TO PARTICIPATE

No ethical approval was required for this study.

### AVAILABILITY OF DATA AND MATERIALS

CLARC is freely available through GitHub (https://github.com/IndraGonz/CLARC) under an MIT license. All code used to generate the analyses in this manuscript can be found at https://github.com/IndraGonz/2024_GonzalezOjeda_CLARC and the output data for the pangenome analyses presented here can be found in the Zenodo folder with DOI: 10.5281/zenodo.14187853 (https://zenodo.org/records/14187853).

### AUTHOR CONTRIBUTIONS

IGO, WPH, and ML conceived the research idea. IGO implemented the pipeline and performed the formal analyses. IGO, SGP, PM, TA, and ML contributed to the data analysis and interpretation. LRG and LLH provided access and insight into the pneumococcal datasets. IGO and ML wrote the first draft of the article. All authors contributed to the manuscript review and editing, approved the final version and agreed to submit it to the journal.

## ACKNOWLEDGEMENTS

The authors acknowledge Katherine L. O’Brien for support in the collection and curation of the pneumococcal datasets. Also, we acknowledge Xueting Qiu and Maximillian Marin for bioinformatics support and fruitful discussions. Lastly, we thank Ruchita Balasubramanian and Léa Cavalli for helpful comments and testing.

## References

1. Marie Touchon, Claire Hoede, Olivier Tenaillon, Valérie Barbe, et al. Organised Genome Dynamics in the Escherichia coli Species Results in Highly Diverse Adaptive Paths. PLOS Genetics, 5(1):e1000344, January 2009. ISSN 1553-7404. doi: 10.1371/journal.pgen.1000344. Publisher: Public Library of Science.

2. Van Rossum, Thea and Ferretti, Pamela and Maistrenko Oleksandr M. and Bork, Peer. Diversity within species: interpreting strains in microbiomes. Nature Reviews Microbiology, 18(9):491–506, September 2020. ISSN 1740-1534. doi: 10.1038/s41579-020-0368-1. Publisher: Nature Publishing Group.

3. Jeroen Geurtsen, Mark de Been, Eveline Weerdenburg, Aldert Zomer, et al. Genomics and pathotypes of the many faces of Escherichia coli. FEMS Microbiology Reviews, 46(6):fuac031, November 2022. ISSN 0168-6445. doi: 10.1093/femsre/fuac031.

4. Arnold, Brian J. and Huang, I.-Ting and Hanage William P. Horizontal gene transfer and adaptive evolution in bacteria. Nature Reviews Microbiology, 20(4):206–218, April 2022. ISSN 1740-1534. doi: 10.1038/s41579-021-00650-4. Publisher: Nature Publishing Group.

5. Alan McNally, Teemu Kallonen, Christopher Connor, Khalil Abudahab, et al. Diversification of Colonization Factors in a Multidrug-Resistant Escherichia coli Lineage Evolving under Negative Frequency-Dependent Selection. mBio, 10(2):10.1128/mbio.00644–19, April 2019. doi: 10.1128/mbio.00644-19. Publisher: American Society for Microbiology.

6. Dhiraj Sinha, Xifeng Sun, Mudra Khare, Michel Drancourt, et al. Pangenome analysis and virulence profiling of Streptococcus intermedius. BMC Genomics, 22(1):522, July 2021. ISSN 1471-2164. doi: 10.1186/s12864-021-07829-2.

7. Fatima Shahid, Tahreem Zaheer, Shifa Tariq Ashraf, Muhammad Shehroz, et al. Chimeric vaccine designs against Acinetobacter baumannii using pan genome and reverse vaccinology approaches. Scientific Reports, 11(1):13213, June 2021. ISSN 2045-2322. doi: 10.1038/s41598-021-92501-8. Publisher: Nature Publishing Group.

8. Hervé Tettelin Vega Masignani, Michael J. Cieslewicz, Claudio Donati, et al. Genome analysis of multiple pathogenic isolates of Streptococcus agalactiae: Implications for the microbial “pan-genome”. Proceedings of the National Academy of Sciences, 102(39):13950–13955, September 2005. doi: 10.1073/pnas.0506758102. Publisher: Proceedings of the National Academy of Sciences.

9. Gabrielle L Harrow, John A Lees, William P Hanage, Marc Lipsitch, et al. Negative frequency-dependent selection and asymmetrical transformation stabilise multi-strain bacterial population structures. The ISME Journal, 15(5):1523–1538, May 2021. ISSN 1751-7362. doi: 10.1038/s41396-020-00867-w.

10. Molari, Marco and Shaw Liam P. and Neher Richard A. Evolutionary dynamics of genome structure and content among closely related bacteria, July 2024. Pages: 2024.07.08.602537 Section: New Results.

11. Andrew J. Page, Carla A. Cummins, Martin Hunt, Vanessa K. Wong, et al. Roary: rapid large-scale prokaryote pan genome analysis. Bioinformatics, 31(22):3691–3693, November 2015. ISSN 1367-4803. doi: 10.1093/bioinformatics/btv421.

12. Ding, Wei and Baumdicker, Franz and Neher Richard A. panX: pan-genome analysis and exploration. Nucleic Acids Research, 46(1):e5, January 2018. ISSN 0305-1048. doi: 10.1093/nar/gkx977.

13. Sion C Bayliss, Harry A Thorpe, Nicola M Coyle, Samuel K Sheppard, et al. PIRATE: A fast and scalable pangenomics toolbox for clustering diverged orthologues in bacteria. GigaScience, 8(10):giz119, October 2019. ISSN 2047-217X. doi: 10.1093/gigascience/giz119.

14. Guillaume Gautreau, Adelme Bazin, Mathieu Gachet, Rémi Planel, et al. PPanGGOLiN: Depicting microbial diversity via a partitioned pangenome graph. PLOS Computational Biology, 16(3):e1007732, March 2020. ISSN 1553-7358. doi: 10.1371/journal.pcbi.1007732. Publisher: Public Library of Science.

15. Gerry Tonkin-Hill, Neil MacAlasdair, Christopher Ruis, Aaron Weimann, et al. Producing polished prokaryotic pangenomes with the Panaroo pipeline. Genome Biology, 21(1):180, July 2020. ISSN 1474-760X. doi: 10.1186/s13059-020-02090-4.

16. Michael Y Galperin, David M Kristensen, Kira S Makarova, Yuri I Wolf, et al. Microbial genome analysis: the COG approach. Briefings in Bioinformatics, 20(4):1063–1070, July 2019. ISSN 1477-4054. doi: 10.1093/bib/bbx117.

17. Lamkiewicz, Kevin and Barf, Lisa-Marie and Sachse, Konrad and Hölzer, Martin. RIBAP: a comprehensive bacterial core genome annotation pipeline for pangenome calculation beyond the species level. Genome Biology, 25(1):170, July 2024. ISSN 1474-760X. doi: 10.1186/s13059-024-03312-9.

18. Maximillian G. Marin, Christoph Wippel, Natalia Quinones-Olvera, Mahboobeh Behruznia, et al. Analysis of the limited M.tuberculosis accessory genome reveals potential pitfalls of pan-genome analysis approaches, March 2024. Pages: 2024.03.21.586149 Section: New Results.

19. Saioa Manzano-Morales, Yang Liu, Sara González-Bodí, Jaime Huerta-Cepas, et al. Comparison of gene clustering criteria reveals intrinsic uncertainty in pangenome analyses. Genome Biology, 24(1):250, October 2023. ISSN 1474-760X. doi: 10.1186/s13059-023-03089-3.

20. Alan McNally, Yaara Oren, Darren Kelly, Ben Pascoe, et al. Combined Analysis of Variation in Core, Accessory and Regulatory Genome Regions Provides a Super-Resolution View into the Evolution of Bacterial Populations. PLOS Genetics, 12(9):e1006280, September 2016. ISSN 1553-7404. doi: 10.1371/journal.pgen.1006280. Publisher: Public Library of Science.

21. Preska Steinberg, Asher and Lin, Mingzhi and Kussell, Edo. Core genes can have higher recombination rates than accessory genes within global microbial populations. eLife, 11:e78533, July 2022. ISSN 2050-084X. doi: 10.7554/eLife.78533. Publisher: eLife Sciences Publications, Ltd.

22. Chrispin Chaguza, Dorota Jamrozy, Merijn W. Bijlsma, Taco W. Kuijpers, et al. Population genomics of Group B Streptococcus reveals the genetics of neonatal disease onset and meningeal invasion. Nature Communications, 13(1):4215, July 2022. ISSN 2041-1723. doi: 10.1038/s41467-022-31858-4. Publisher: Nature Publishing Group.

23. Rebecca A. Gladstone, Stephanie W. Lo, John A. Lees, Nicholas J. Croucher, et al. International genomic definition of pneumococcal lineages, to contextualise disease, antibiotic resistance and vaccine impact. eBioMedicine, 43:338–346, May 2019. ISSN 2352-3964. doi: 10.1016/j.ebiom.2019.04.021. Publisher: Elsevier.

24. Brian D. Ondov, Todd J. Treangen, Páll Melsted, Adam B. Mallonee, et al. Mash: fast genome and metagenome distance estimation using MinHash. Genome Biology, 17(1):132, June 2016. ISSN 1474-760X. doi: 10.1186/s13059-016-0997-x.

25. Domingo-Sananes, Maria Rosa and McInerney James O. Mechanisms That Shape Microbial Pangenomes. Trends in Microbiology, 29(6):493–503, June 2021. ISSN 0966-842X, 1878-4380. doi: 10.1016/j.tim.2020.12.004. Publisher: Elsevier.

26. Claudio Donati, N. Luisa Hiller, Hervé Tettelin Alessandro Muzzi, et al. Structure and dynamics of the pan-genome of Streptococcus pneumoniae and closely related species. Genome Biology, 11(10):R107, October 2010. ISSN 1474-760X. doi: 10.1186/gb-2010-11-10-r107.

27. Alejandro Couce, Anurag Limdi, Melanie Magnan, Siân V. Owen, et al. Changing fitness effects of mutations through long-term bacterial evolution. Science, 383(6681):eadd1417, January 2024. doi: 10.1126/science.add1417. Publisher: American Association for the Advancement of Science.

28. van Opijnen, Tim and Bodi Kip L. and Camilli, Andrew. Tn-seq: high-throughput parallel sequencing for fitness and genetic interaction studies in microorganisms. Nature Methods, 6 (10):767–772, October 2009. ISSN 1548-7105. doi: 10.1038/nmeth.1377. Publisher: Nature Publishing Group.

29. Segata, Nicola and Huttenhower, Curtis. Toward an Efficient Method of Identifying Core Genes for Evolutionary and Functional Microbial Phylogenies. PLOS ONE, 6(9):e24704, September 2011. ISSN 1932-6203. doi: 10.1371/journal.pone.0024704. Publisher: Public Library of Science.

30. Jukka Corander, Christophe Fraser, Michael U. Gutmann, Brian Arnold, et al. Frequencydependent selection in vaccine-associated pneumococcal population dynamics. Nature Ecology & Evolution, 1(12):1950–1960, December 2017. ISSN 2397-334X. doi: 10.1038/s41559-017-0337-x. Publisher: Nature Publishing Group.

31. Taj Azarian, Pamela P. Martinez, Brian J. Arnold, Xueting Qiu, et al. Frequency-dependent selection can forecast evolution in Streptococcus pneumoniae. PLOS Biology, 18(10):e3000878, October 2020. ISSN 1545-7885. doi: 10.1371/journal.pbio.3000878. Publisher: Public Library of Science.

32. Yildirim, Inci and Shea Kimberly M. and Pelton Stephen I. Pneumococcal Disease in the Era of Pneumococcal Conjugate Vaccine. Infectious Disease Clinics, 29(4):679–697, December 2015. ISSN 0891-5520, 1557-9824. doi: 10.1016/j.idc.2015.07.009. Publisher: Elsevier.

33. Christie, Mark R. and McNickle Gordon G. Negative frequency dependent selection unites ecology and evolution. Ecology and Evolution, 13(7):e10327, 2023. ISSN 2045-7758. doi: 10.1002/ece3.10327. _eprint: https://onlinelibrary.wiley.com/doi/pdf/10.1002/ece3.10327.

34. Li, Weizhong and Godzik, Adam. Cd-hit: a fast program for clustering and comparing large sets of protein or nucleotide sequences. Bioinformatics, 22(13):1658–1659, July 2006. ISSN 1367-4803. doi: 10.1093/bioinformatics/btl158.

35. Enright, A. J. and Dongen, S.Van and Ouzounis, C.A. An efficient algorithm for largescale detection of protein families. Nucleic Acids Research, 30(7):1575, April 2002. doi: 10.1093/nar/30.7.1575.

36. Horsfield, Samuel T. and Tonkin-Hill, Gerry and Croucher Nicholas J. and Lees, John Accurate and fast graph-based pangenome annotation and clustering with ggCaller. Genome Research, page gr.277733.123, August 2023. ISSN 1088-9051, 1549-5469. doi: 10.1101/gr.277733.123. Company: Cold Spring Harbor Laboratory Press Distributor: Cold Spring Harbor Laboratory Press Institution: Cold Spring Harbor Laboratory Press Label: Cold Spring Harbor Laboratory Press Publisher: Cold Spring Harbor Lab.

37. Ponting, Chris P. Issues in predicting protein function from sequence. Briefings in Bioinformatics, 2(1):19–29, March 2001. ISSN 1467-5463. doi: 10.1093/bib/2.1.19.

38. Gabaldón, Toni and Koonin Eugene V. Functional and evolutionary implications of gene orthology. Nature Reviews Genetics, 14(5):360–366, May 2013. ISSN 1471-0064. doi: 10.1038/nrg3456. Publisher: Nature Publishing Group.

39. Tonkin-Hill, Gerry and Corander, Jukka and Parkhill, Julian. Challenges in prokaryote pangenomics. Microbial Genomics, 9(5):001021, 2023. ISSN 2057-5858. doi: 10.1099/mgen.0.001021. Publisher: Microbiology Society,.

40. Athina Gavriilidou, Emilian Paulitz, Christian Resl, Nadine Ziemert, et al. Goldfinder: Unraveling Networks of Gene Co-occurrence and Avoidance in Bacterial Pangenomes, May 2024. Pages: 2024.04.29.591652 Section: New Results.

41. William P. Hanage, Cynthia J. Bishop, Susan S. Huang, Abbie E. Stevenson, et al. Carried Pneumococci in Massachusetts Children: The Contribution of Clonal Expansion and Serotype Switching. The Pediatric Infectious Disease Journal, 30(4):302, April 2011. ISSN 0891-3668. doi: 10.1097/INF.0b013e318201a154.

42. Patrick K. Mitchell, Taj Azarian, Nicholas J. Croucher, Alanna Callendrello, et al. Population genomics of pneumococcal carriage in Massachusetts children following introduction of PCV-13. Microbial Genomics, 5(2):e000252, February 2019. doi: 10.1099/mgen.0.000252.

43. Eugene V. Millar, Katherine L. O’Brien, Elizabeth R. Zell, Melinda A. Bronsdon, et al. Nasopharyngeal Carriage of Streptococcus pneumoniae in Navajo and White Mountain Apache Children Before the Introduction of Pneumococcal Conjugate Vaccine. The Pediatric Infectious Disease Journal, 28(8):711, August 2009. ISSN 0891-3668. doi: 10.1097/INF.0b013e3181a06303.

44. Lindsay R. Grant, Laura L. Hammitt, Sarah E. O’Brien, Michael R. Jacobs, et al. Impact of the 13-Valent Pneumococcal Conjugate Vaccine on Pneumococcal Carriage Among American Indians. The Pediatric Infectious Disease Journal, 35(8):907, August 2016. ISSN 0891-3668. doi: 10.1097/INF.0000000000001207.

45. Katherine L O’Brien, Lawrence H Moulton, Raymond Reid, Robert Weatherholtz, et al. Efficacy and safety of seven-valent conjugate pneumococcal vaccine in American Indian children: group randomised trial. The Lancet, 362(9381):355–361, August 2003. ISSN 0140-6736. doi: 10.1016/S0140-6736(03)14022-6.

46. Claire Chewapreecha, Simon R. Harris, Nicholas J. Croucher, Claudia Turner, et al. Dense genomic sampling identifies highways of pneumococcal recombination. Nature Genetics, 46(3):305–309, March 2014. ISSN 1546-1718. doi: 10.1038/ng.2895. Publisher: Nature Publishing Group.

47. Rebecca A. Gladstone, Johanna M. Jefferies, Anna S. Tocheva, Kate R. Beard, et al. Five winters of pneumococcal serotype replacement in UK carriage following PCV introduction. Vaccine, 33(17):2015–2021, April 2015. ISSN 0264-410X. doi: 10.1016/j.vaccine.2015.03.012.

48. Andries J. van Tonder, James E. Bray, Lucy Roalfe, Rebecca White, et al. Genomics Reveals the Worldwide Distribution of Multidrug-Resistant Serotype 6E Pneumococci. Journal of Clinical Microbiology, 53(7):2271, June 2015. doi: 10.1128/JCM.00744-15.

49. Ellen Heinsbroek, Terence Tafatatha, Amos Phiri, Todd D Swarthout, et al. Pneumococcal carriage in households in Karonga District, Malawi, before and after introduction of 13-valent pneumococcal conjugate vaccination. Vaccine, 36(48):7369–7376, November 2018. ISSN 0264-410X. doi: 10.1016/j.vaccine.2018.10.021.

50. Susan A. Nzenze, Anne von Gottberg, Tinevimbo Shiri, Nadia van Niekerk, et al. Temporal Changes in Pneumococcal Colonization in HIV-infected and HIV-uninfected Mother-Child Pairs Following Transitioning From 7-valent to 13-valent Pneumococcal Conjugate Vaccine, Soweto, South Africa. The Journal of Infectious Diseases, 212(7):1082–1092, October 2015. ISSN 0022-1899. doi: 10.1093/infdis/jiv167.

51. Xueting Qiu, Lesley McGee, Laura L Hammitt, Lindsay R. Grant, et al. Prediction of post-PCV13 pneumococcal evolution using invasive disease data enhanced by inverse-invasiveness weighting. mBio, 15(10):e03355–23, August 2024. doi: 10.1128/mbio.03355-23. Publisher: American Society for Microbiology.

52. Andrey Prjibelski, Dmitry Antipov, Dmitry Meleshko, Alla Lapidus, et al. Using SPAdes De Novo Assembler. Current Protocols in Bioinformatics, 70(1):e102, 2020. ISSN 1934-340X. doi: 10.1002/cpbi.102. _eprint: https://currentprotocols.onlinelibrary.wiley.com/doi/pdf/10.1002/cpbi.102.

53. Seemann, Torsten. Prokka: rapid prokaryotic genome annotation. Bioinformatics, 30(14):2068–2069, July 2014. ISSN 1367-4803. doi: 10.1093/bioinformatics/btu153.

54. Wick, Ryan R. and Judd Louise M. and Gorrie Claire L. and Holt Kathryn E. Unicycler: Resolving bacterial genome assemblies from short and long sequencing reads. PLOS Computational Biology, 13(6):e1005595, June 2017. ISSN 1553-7358. doi: 10.1371/journal.pcbi.1005595. Publisher: Public Library of Science.

55. Gurevich, Alexey and Saveliev, Vladislav and Vyahhi, Nikolay and Tesler, Glenn. QUAST: quality assessment tool for genome assemblies. Bioinformatics, 29(8):1072–1075, April 2013. ISSN 1367-4803. doi: 10.1093/bioinformatics/btt086.

56. John A. Lees, Simon R. Harris, Gerry Tonkin-Hill, Rebecca A. Gladstone, et al. Fast and flexible bacterial genomic epidemiology with PopPUNK. Genome Research, 29(2):304, February 2019. doi: 10.1101/gr.241455.118.

57. Stephen F. Altschul, Warren Gish, Webb Miller, Eugene W. Myers, et al. Basic local alignment search tool. Journal of Molecular Biology, 215(3):403–410, October 1990. ISSN 0022-2836. doi: 10.1016/S0022-2836(05)80360-2.

58. Tatusov, Roman L. and Galperin Michael Y. and Natale Darren A. and Koonin Eugene V. The COG database: a tool for genome-scale analysis of protein functions and evolution. Nucleic Acids Research, 28(1):33–36, January 2000. ISSN 0305-1048. doi: 10.1093/nar/28.1.33.

59. Jaime Huerta-Cepas, Damian Szklarczyk, Davide Heller, Ana Hernández-Plaza, et al. eggNOG 5.0: a hierarchical, functionally and phylogenetically annotated orthology resource based on 5090 organisms and 2502 viruses. Nucleic Acids Research, 47(D1):D309–D314, January 2019. ISSN 0305-1048. doi: 10.1093/nar/gky1085.

60. Brynildsrud, Ola and Bohlin, Jon and Scheffer, Lonneke and Eldholm, Vegard. Rapid scoring of genes in microbial pan-genome-wide association studies with Scoary. Genome Biology, 17(1):238, November 2016. ISSN 1474-760X. doi: 10.1186/s13059-016-1108-8.

61. Carlos P Cantalapiedra, Ana Hernández-Plaza, Ivica Letunic, Peer Bork, et al. eggNOGmapper v2: Functional Annotation, Orthology Assignments, and Domain Prediction at the Metagenomic Scale. Molecular Biology and Evolution, 38(12):5825–5829, December 2021. ISSN 1537-1719. doi: 10.1093/molbev/msab293.

62. Frank, Marguerite and Wolfe, Philip. An algorithm for quadratic programming. Naval Research Logistics Quarterly, 3(1-2):95–110, 1956. ISSN 1931-9193. doi: 10.1002/nav.3800030109. _eprint: https://onlinelibrary.wiley.com/doi/pdf/10.1002/nav.3800030109.

