## Supplementary figure S1, Supplementary figure S2, Supplementary tables S1-S2 for "Linkage-based ortholog refinement in bacterial pangenomes with CLARC"

**ADDITIONAL FILE 1**

**SUPPLEMENTARY TABLES**

**Table S1.** CLARC clusters found in pangenomes generated from the Southwest, USA dataset + 9 closed references (946 genomes) using different tools and parameters.

| **Pangenome tool/parameters** | **Number of accessory CLARC clusters** | **Number of core CLARC clusters** | **Total # of CLARC clusters** |
| --- | --- | --- | --- |
| Roary i98 | 269 | 139 | **408** |
| Roary, i95 (default) | 99 | 21 | **120** |
| Roary, i90 | 95 | 24 | **119** |
| Roary, i80 | 89 | 27 | **116** |
| Panaroo, i98 – strict (default) | 8 | 1 | **9** |
| Panaroo, i98 – moderate | 8 | 2 | **10** |
| Panaroo, i98 – sensitive | 9 | 2 | **11** |
| Panaroo, i95 - strict | 3 | 1 | **4** |
| Panaroo, i95 - moderate | 7 | 1 | **8** |
| Panaroo i95 - sensitive | 7 | 1 | **8** |
| Panaroo, i90 – strict | 2 | 1 | **3** |
| Panaroo, i90 – moderate | 3 | 2 | **5** |
| Panaroo, i90 – sensitive | 3 | 2 | **5** |
| Panaroo, i80 – strict | 2 | 0 | **2** |
| Panaroo, i80 – moderate | 3 | 1 | **4** |
| Panaroo, i80 – sensitive | 3 | 1 | **4** |

**Table S2.** CLARC clusters found in pangenomes generated from the samples in all carriage datasets + 9 closed references (8907 genomes) using different tools and parameters.

| **Pangenome tool/parameters** | **Number of accessory CLARC clusters** | **Number of core CLARC clusters** | **Total # of CLARC clusters** |
| --- | --- | --- | --- |
| Roary, i98 | 401 | 479 | **880** |
| Roary, i95 (default) | 274 | 453 | **727** |
| Roary, i90 | 247 | 452 | **699** |
| Roary, i80 | 217 | 430 | **647** |
| Panaroo, i98 – strict (default) | 2 | 0 | **2** |
| Panaroo, i98 – moderate | 3 | 0 | **3** |
| Panaroo, i98 – sensitive | 5 | 0 | **5** |
| Panaroo, i95 - strict | 2 | 0 | **2** |
| Panaroo, i95 - moderate | 6 | 0 | **6** |
| Panaroo, i95 - sensitive | 1 | 0 | **1** |
| Panaroo, i90 – strict | 1 | 0 | **1** |
| Panaroo, i90 – moderate | 3 | 0 | **3** |
| Panaroo, i90 – sensitive | 0 | 0 | **0** |
| Panaroo, i80 – strict | 2 | 0 | **2** |
| Panaroo, i80 – moderate | 2 | 1 | **3** |
| Panaroo, i80 – sensitive | 1 | 1 | **2** |

**SUPPLEMENTARY FIGURES**

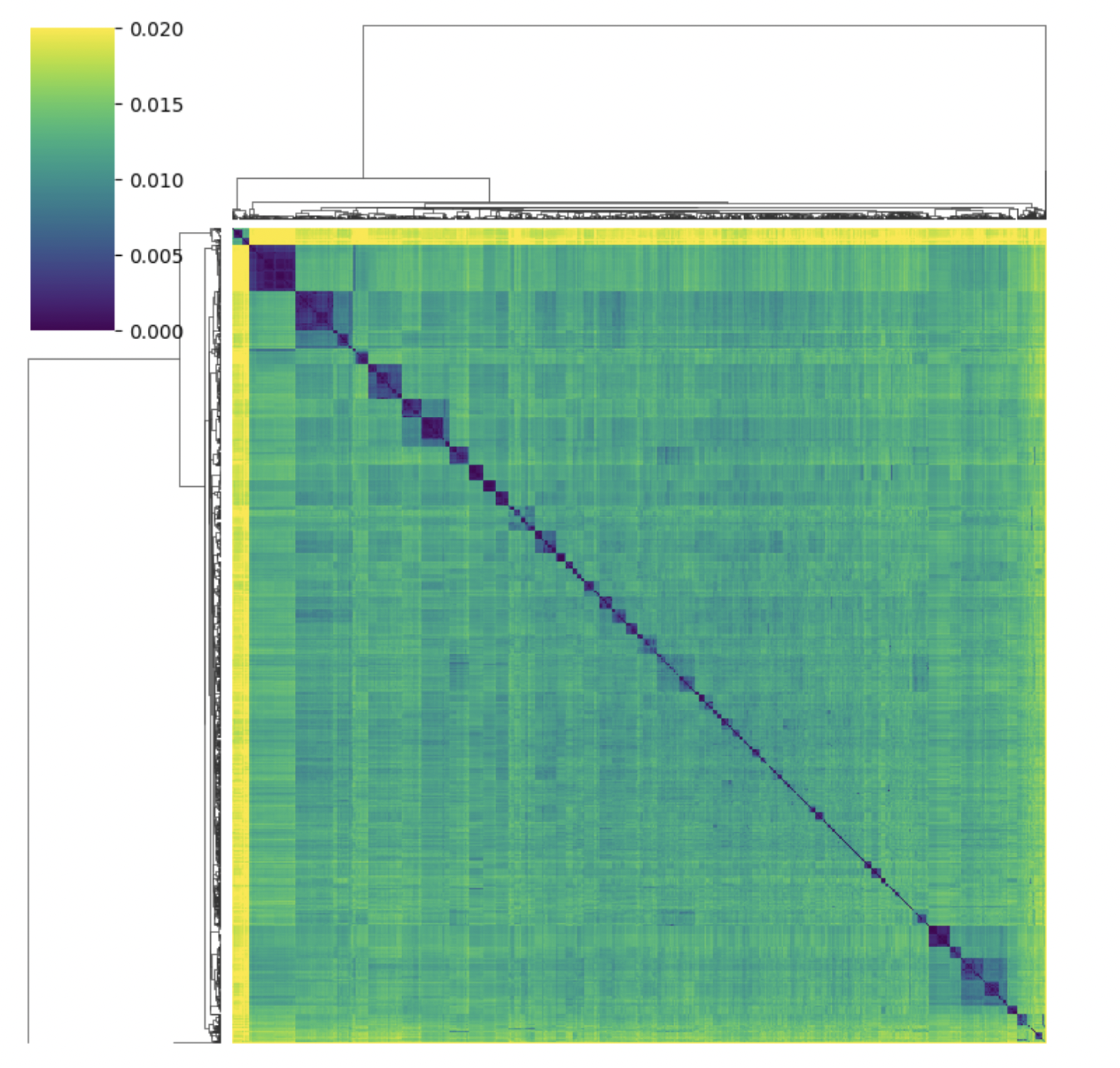

**Figure S1.** *Mash distances for all 8,907 S. pneumoniae genomes used in this study.* Average genomic distance is 1.24%. Mash was run on default parameters.

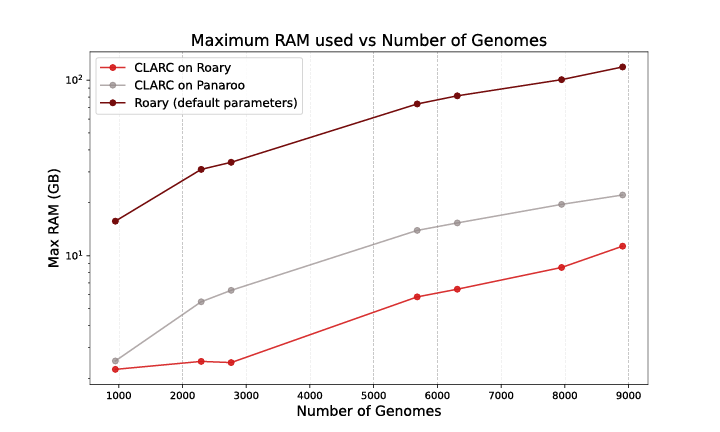

**Figure S2.** *Computational performance of CLARC.* Performance for Panaroo is not pictured because running Panaroo on all pneumococcal genomes in one job required more computational resources than available in the FASRC computing cluster. All multi-dataset Panaroo analyses shown in the main text were obtained through a 2-step process where we ran Panaroo individually on each carriage dataset, and then ran the appropriate merge commands within Panaroo to obtain the pangenome definitions of runs with >1,000 genomes.
